# Automated sample preparation for high-throughput single-cell proteomics

**DOI:** 10.1101/399774

**Authors:** Harrison Specht, Guillaume Harmange, David H. Perlman, Edward Emmott, Zachary Niziolek, Bogdan Budnik, Nikolai Slavov

## Abstract

A major limitation to applying quantitative LC-MS/MS proteomics to small samples, such as single cells, are the losses incured during sample cleanup. To relieve this limitation, we developed a Minimal ProteOmic sample Preparation (mPOP) method for culture-grown mammalian cells. mPOP obviates cleanup and thus eliminates cleanup-related losses while expediting sample preparation and simplifying its automation. Bulk SILAC samples processed by mPOP or by conventional urea-based methods indicated that mPOP results in complete cell lysis and accurate relative quantification. We integrated mPOP lysis with the Single Cell ProtEomics by Mass Spectrometry (SCoPE-MS) sample preparation, and benchmarked the quantification of such samples on a Q-exactive instrument. The results demonstrate low noise and high technical reproducibility. Then, we FACS sorted single U-937, HEK-293, and mouse ES cells into 96-well plates and analyzed them by automated mPOP and SCoPE-MS. The quantified proteins enabled separating the single cells by cell-type and cell-division-cycle phase.

## Introduction

Methods for preparing sub-microgram protein samples for LC-MS/MS often use sophisticated custom-made equipment^1–3^ and generally lyse cells by detergents or chaotropic agents like urea^3,4^. Techniques using these chemicals are robust but require that the chaotropic agents or detergents be removed before MS analysis since these chemicals are incompatible with MS^4^. Some cleanup methods, such as SP3^5^ and iST^6^ perform very well even for microgram samples^4,7^. Yet losses are more significant for the preparation of low-abundance samples, such as single cells. Further more, cleanup steps complicate automation and may introduce variability between samples. Thus, avoiding cleanup stages can reduce losses while increasing throughput and consistency^8^. A cell lysis method that does not require MS-incompatible chemicals and thus can be used for LC-MS/MS without cleaning is focused acoustic sonication (FAS)^4,9^. We successfully used FAS to obviate cleaning up single-cell lysates and to develop Single Cell ProtEomics by Mass Spectrometry (SCoPE-MS)^10^. While FAS resulted in clean lysis, it required significant volumes (5 *-* 10*µl*), was low-throughput, and used expensive consumables and equipment^10^. These limitations hinder its potential for high-throughput single-cell proteomics^8^.

## Results

To relieve these limitations, we sought to develop a method for lysing cells in pure water that is high-throughput, inexpensive, easily-automated, compatible with small lysis volumes and only uses common, inexpensive, commercial laboratory equipment. We started by evaluating the LC-MS/MS compatibility of lysis methods developed for other applications and found that protein extraction was rather incomplete compared to methods validated for LC-MS/MS that use detergents and chaotropic chemicals. Among these methods, freeze-thaw cycles in pure water showed the most promise, and we iteratively optimized it to increase its robustness and the efficiency of delivering peptides for LC-MS/MS analysis while preserving the physiological state of the analyzed cells. These efforts culminated in mPOP, a method that lyses culture-grown mammalian cells by a freeze-heat cycle (*-*80^*°*^*C* to 90^*°*^*C*) in small droplets of pure water, as illustrated in Fig. 1a.

**Figure 1.**
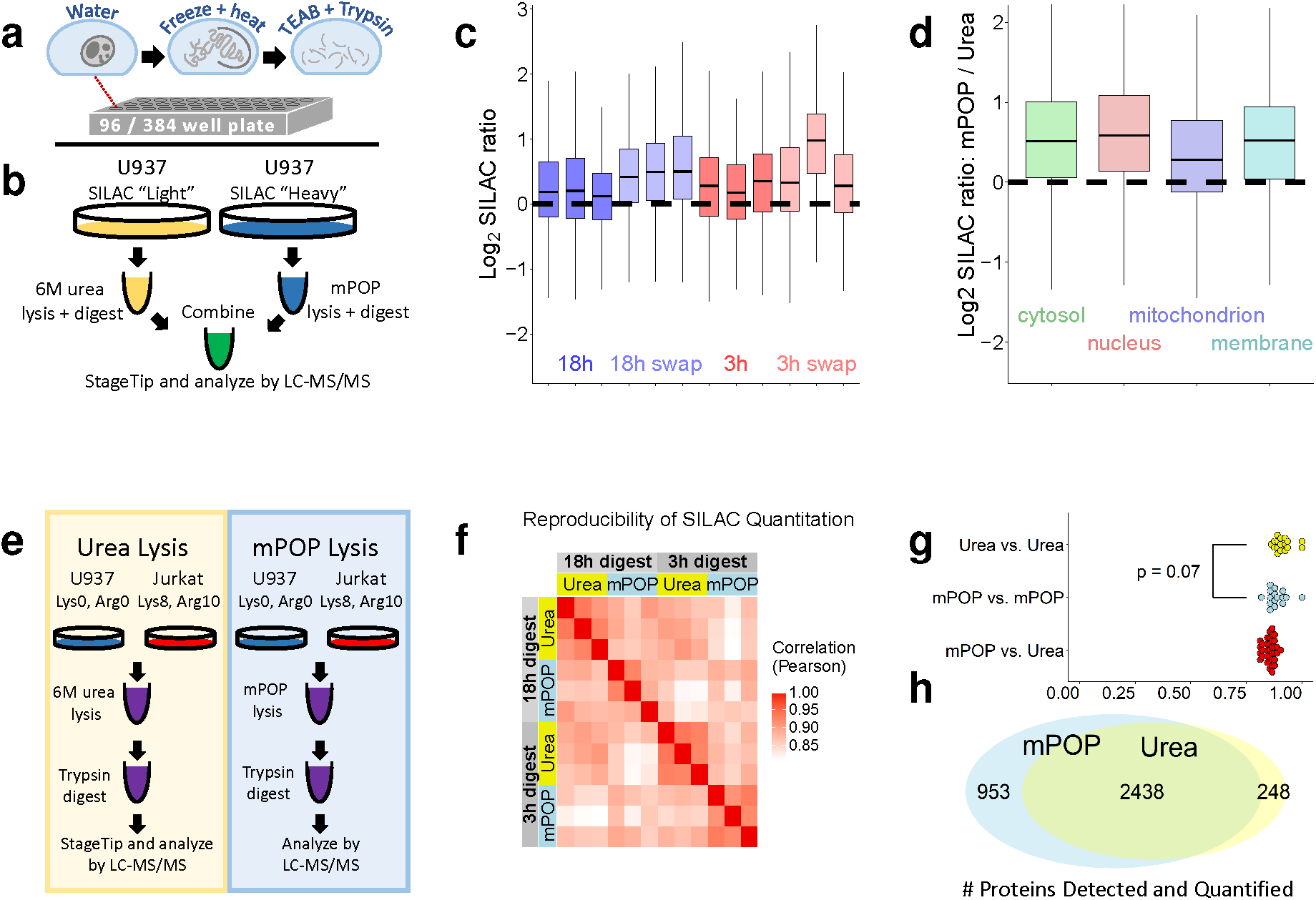
Validating mPOP cell lysis by comparison to urea lysis using SILAC labeling. (**a**) Conceptual diagram of a high-throughput mPOP workflow: cells are lysed by a freeze-heat cycle (-80°C to 90°C), pH adjusted by triethylammonium bicarbonate (TEAB) and enzymatically digested to peptides. (**b**) Schematic of experiments comparing lysis yield: Cells were sorted by FACS, lysed by either mPOP or 6M urea, and the proteins digested to peptides by trypsin. Lysates were combined, cleaned from urea by StageTip, concentrated, and analyzed by LC-MS/MS. (**c**) Lysis by mPOP compared to 6M urea across 12 replicates, including SILAC label swaps and two digestion conditions, 3 hours and 18 hours. Lysis efficiency was quantified by the distribution of mPOP / Urea peptide SILAC ratios, with equivalent lysis and digestion displayed with the dotted line at zero. (**d**) SILAC ratios from panel (c) grouped by cellular compartment indicate that mPOP efficiently extracts proteins from all compartments. (**e**) Schematic of comparing quantification by in cells lysed by urea or mPOP. Cells were sorted by FACS into tubes to contain 10,000 ”heavy” Jurkat cells and ”light” U-937 cells, lysed by either mPOP or 6M urea, the proteins digested to peptides by trypsin, urea removed by StageTip clean-up if necessary, concentrated, and analyzed by LC-MS/MS. (**f**) Correlation matrix of all biological replicates produced from the experiment described in panel (d) including two digestion conditions: 3 hours and 18 hours. (**g**) Correlations from panel (e) displayed as distributions. Differences between mPOP and urea are insignificant, p-value = 0.07, based on Kolmogorov-Smirnov test. (**h**) Proteins identified and quantified by mPOP and urea lysis overlap significantly, but mPOP identified 953 proteins not identified in the urea lysates.

For bottom-up proteomics, the cell lysate is then digested in 10 *ng/µl* of the protease trypsin. This simple procedure allowed us to design a proteomics sample preparation using only a MS-compatible digestion buffer (Triethylammonium bicarbonate, pH 8.0), trypsin, and formic acid. Crucially, mPOP allows minimizing volumes, which reduces sample losses and reagents used. It also allows sample preparation in 96/384 well-plates, which enabled simultaneous processing of many samples in parallel. Furthermore, the obviation of cleanup allowed us to easily automate mPOP sample preparation with inexpensive PCR thermocyclers and liquid dispensers.

### Evaluating the completeness of lysis and protein extraction

We sought to directly compare the lysis efficiency of mPOP to that of standard 6M urea lysis using the experimental design in Fig. 1b. Urea was chosen because it is a widely-used lysis method for LC-MS/MS that compares favorably to other methods and its accessibility facilitates replication^11^. We lysed samples of FACS sorted U-937 cells with either mPOP or urea, Fig. 1b. Each sample was comprised of 10,000 cells having either light SILAC or heavy SILAC label. Samples of 10,000 cells were chosen to provide enough proteins so that clean-up losses by StageTip^12^ (which is required by urea lysis) are affordable and lysis efficiency can be evaluated independently from cleanup-losses. Light cells lysed by urea were mixed with heavy cells lysed by mPOP, Fig. 1b. To control for possible biases, we also performed a label swap in which heavy cells lysed by urea were mixed with light cells lysed by mPOP. The mixtures of light and heavy cell-lysates were cleaned-up by StageTip to remove urea. This design incurred unnecessary clean-up losses from the mPOP lysates (since they do not need to be cleaned), but it allowed us to evaluate the lysis efficiency of mPOP to that of 6M urea independently of cleanup losses since the cleanup losses in this experiment occur after the mixing and are identical for both lysis methods. These samples were analyzed by LC-MS/MS, and the relative abundance of each peptide between the heavy and light lysates quantified with its SILAC ratio. The distributions of SILAC ratios for all peptides (Fig. 1c) indicate that most peptides have higher abundances in samples lysed by mPOP, suggesting that mPOP allows delivering peptides to MS analysis at least as efficiently as urea lysis. To examine potential bias in the extraction of proteins, we analyzed the distribution of SILAC ratios partitioned by cellular compartment, including both compartments expected to be difficult and easy to lyse, Fig. 1c. The results indicate that mPOP lysis outperforms urea lysis for proteins residing in the cytosol, mitochondrion, nucleus, and the cell membrane. Indeed, no gene sets with greater than two unique proteins favor urea lysis over mPOP.

### Evaluating quantification accuracy

Having established that mPOP lyses cells efficiently, we sought to evaluate the reproducibility of relative protein quantification between mPOP and urea lysis using the experimental design outlined in Fig. 1e. FACS-sorted samples of 10,000 heavy SILAC Jurkat cells were combined with 10,000 light SILAC U937 cells in the same tube. These sample were lysed with either mPOP or urea. All cell-lysates were digested by trypsin (urea-containing samples after dilution to *<* 1*M*) for either 3 or 18 hours. Then, trypsin was quenched with 1% by volume formic acid, Fig. 1e. To remove the urea, samples were cleaned-up by StageTip. All samples were analyzed on a Orbitrap Lumos, and the relative protein levels between Jurkat and U937 cells estimated with the corresponding SILAC ratios computed by MaxQuant. To compare the consistency of quantification within and between lysis methods, we compared the pairwise correlations between the SILAC ratios of all samples, Fig. 1ef. The correlations ranged between 0.78 and 1, indicating excellent reproducibility both within and across lysis methods. Furthermore, we found the mean coefficient of variation of peptide SILAC ratios to be *<* 10% for both mPOP and urea replicates, Fig. S1. Since we did not mix the samples lysed by mPOP and by urea, we could compare proteome coverage between the two methods (Fig. 1h). The number of proteins identified and quantified using urea lysis is comparable to that from similar label-free studies^4^. Almost all of these proteins were identified and quantified by mPOP as well (2,438 proteins), but mPOP samples contained an additional 953 proteins, Fig. 1h.

### Combining mPOP and SCoPE-MS

Our results with 10,000 cells demonstrate that mPOP performs as well or better than urea lysis in terms of (*i*) efficiency of proteome extraction (Fig. 1a-d), (*ii*) quantification accuracy (Fig. 1e-g), and (iii) depth of proteome coverage (Fig. 1f). Next, we turn to the key advantages of mPOP, namely parallel and automated preparation of samples that are too small to be cleaned-up without significant losses. To further reduce losses during nano liquid chromatography (nLC), enhance sequence identification, and increase throughput, we used the carrier design that we introduced with SCoPE-MS^10^ but lysed the cells with mPOP instead by FAS. Introducing mPOP allowed us to reduce lysis volumes 10-fold, from 10*µl* to 1*µl*, to reduce the cost of consumables and equipment over 100-fold, and to increase throughput of sample preparation over 100-fold by preparing many samples in parallel.

### Benchmarking instrument noise with SCoPE-MS design

Before applying mPOP to prepare and analyze single-cell proteomes, we sought to estimate the instrument measurement noise in the context of SCoPE-MS sets. This estimate is motivated by our concern that factors unique to ultra-low abundance samples, such as counting noise^3,8^, may undermine measurement accuracy. To isolate the noise in instrument (Q-exactive) measurement from noise due to biological variation and sample preparation, we used mPOP to prepare a 100*×M* SCoPE-MS sample with two carrier channels (126C-Jurkat cells; 127N-U-937 cells) and 6 interleaved single-cell channels (3 Jurkat and 3 U-937 cells), as shown in **Supplementary Fig. 1a**. Thus 1% dilution (1*×M*) represented the protein abundances expected for single-cell SCoPE-MS set; see **Supplementary Fig. 1a**. Although we did not clean the sample, the 1*×M* dilutions were clean enough to be analyzed by direct injection using a commercial Waters column, and resulted in robust, ion-rich spectra, Fig. 2a. Each 1*×M* injection was analyzed for only 60 min since our goal was to optimize the number of proteins quantified across many cells, rather then the number of proteins quantified per injection^8^. Indeed, we find that the number of peptides quantified across many cells, and thus suitable for biological analysis, increases with the number of analyzed cells, Fig. 2b. The number of confidently identified proteins can be increased up to 50% by applying DART-ID, a data-driven Bayesian framework that uses retention time evidence to enhance peptide sequence identification^13^. Taken together, these results suggest that mPOP supports the preparation of SCoPE-MS sets from low-input samples.

**Figure 2.**
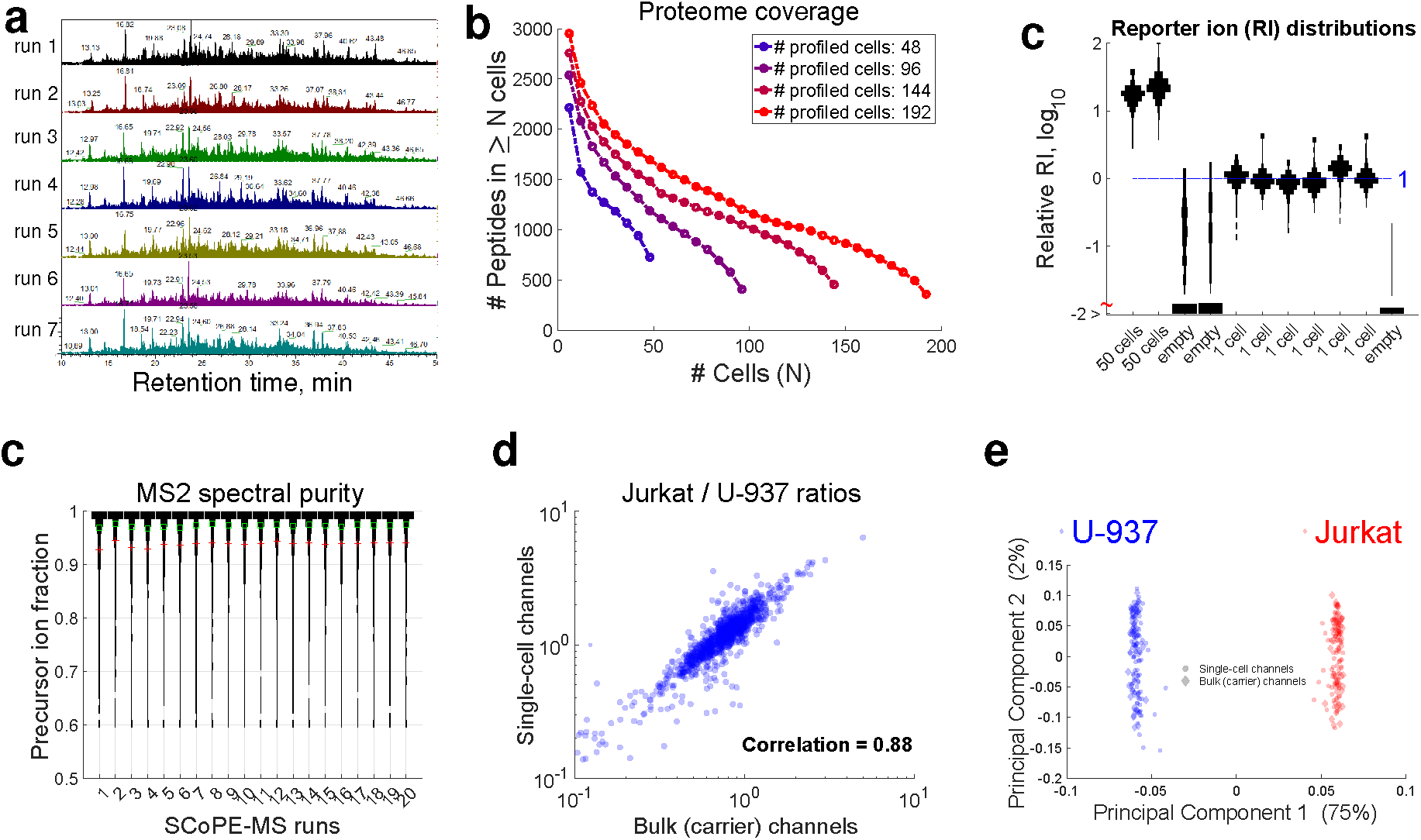
Benchmarking diluted SCoPE-MS sets prepared by mPOP. (**a**) We lysed FACS sorted Jurkat and U-937 cells with mPOP, and prepared a 1 *× M* SCoPE-MS set as described in **Supplementary Fig. 1**a. Direct injections of 1 *×M* SCoPE-MS set on a commercial Waters column resulted in reproducible and rich spectra. The y-axes of all panels range from 0 to 3*×* 10^8^. (**b**) Proteome coverage increases with the number of quantified cells. All identifications are based on spectra only, not using retention times. (**c**) The reporter ion (RI) signal in single-cell channels is much larger than in the empty channels. (**d**) mPOP and SCoPE-MS allow for pure MS2 spectra. (**e**) Relative peptide levels estimated from single-cell SCoPE-MS channels are very similar to the corresponding estimates from the carrier (bluk) channels. (**f**) Principal component analysis separates perfectly single-cell and carrier channels dependent on whether they correspond to Jurkat or to U-937 cells. All quantified proteins were used for this analysis and each protein was normalized separately to a mean levels of one for the carrier channels and the single-cell channels. Without this normalization, PC2 separated the carrier from the single-cell channels, **Supplementary Fig. 1**b.

Next we benchmarked the signal to noise ratio (SNR) and the relative quantification from the single-cell channels in the 1 *× M* samples. To evaluate the SNR, we compared the distributions of relative reporter (RI) ion ratios from single-cell channels and for empty channels, Fig. 2c. We found that the majority of the peptides have orders of magnitude lower signal in the empty channels compared to the single-cell channels, despite the low level of isotopic contamination from the carrier channels, Fig. 2c. This observation and the high purity of the MS2 spectra shown in Fig. 2d suggest that the single-cell RIs contain peptide signal. To evaluate whether this signal is quantitative, we benchmarked the Jurkat / U-937 ratios estimated from single-cell channels against the corresponding ratios estimated from the carrier channels, Fig. 2e. The high concordance of these estimates (Spearman *ρ* = 0.88) strongly indicate that the instrument (Q-exactive) noise in quantifying single-cell-level peptides as part of the SCoPE-MS samples is small, consistent with our arguments that the abundance of proteins in mammalian single cells is high-enough to minimize the sampling (counting) noise^8^. To further evaluate relative quantification, beyond the results for a single SCoPE-MS set displayed in Fig. 2e, we consolidated the data from 34 SCoPE-MS sets and computed all pairwise correlations among single-cell and carrier channels. This 272-dimensional matrix was projected just on its first two principal components (PC). When the carrier and single-cell channels are normalized, PC1 separates perfectly all channels corresponding to Jurkat or U-937 cells, accounting for the majority of the variance (75%) in the data. Without the normalization, PC1 still perfectly separates the measurements by cell type, and PC2 separates the single-cell channels from the carrier channels; see **Supplementary Fig. 1b**. Crucially, the single-cell channels separate the same way as the carrier channels, indicating that all single-cell channels were correctly quantified in our work-flow.

### Quantifying single cell proteomes

Having demonstrated that 1*×M* sets can be analyzed with low noise by LC-MS/MS on Q-exactive, we next applied mPOP to the analysis of single cells that were FACS sorted into 96-well plates, one cell per well, Fig. 3a. Unlike the results from Fig. 2 that characterize just technical variability, this analysis of single cells includes additional variability due to the handing of single cells and due to biological differences between single cells. As a first proof of principle, we again sorted HEK-293 and U-937 cells, and found that when processed by mPOP and SCoPE-MS, their proteomes separate along the first principal component of PCA analysis, Fig. 3b.

**Figure 3.**
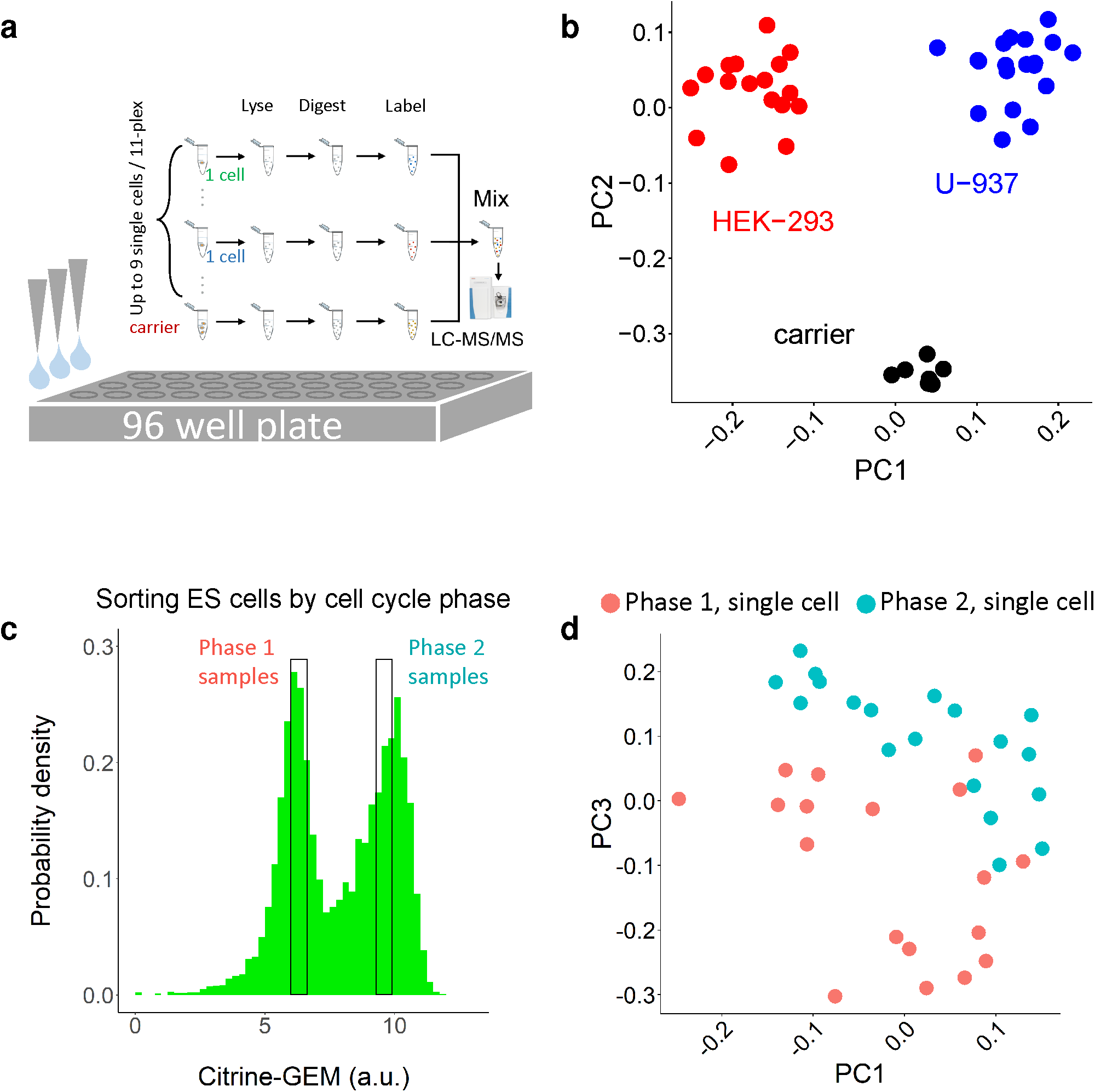
mPOP enables proteomic analysis of cancer cell lines and the mouse embryonic cell cycle from asynchronous single cells. (**a**) Experimental design for high-throughput, low-input proteomics with mPOP combined with SCoPE-MS. Single cells are sorted into 96-well plates and used for SCoPE-MS sets. While these sets can include up to nine single cells / set, the data shown in panels b and c used seven and six single cells / set respectively, because some TMT channels were used for controls; see Methods. (**b**) A principal component analysis (PCA) of single HEK-293 and U-937 cells. The cells were sorted by FACS in a 96-well plate, one cell per well, and processed by mPOP and SCoPE-MS. The first principal component (PC1) separates the projected single-cell proteomes by cell type. (**c**) Mouse embryonic stem cells expressing the FUCCI system^14^ were sorted by Aria FACS into a 96-well plate, one cell per well, from two distinct phases of the cell division cycle. The phases were inferred from the fluorescent protein Citrine, fused to partial sequences of Geminin that is a ubiquitin-targe1t6of the anaphase promoting complex. (d) The proteins with the largest variance separate the single cells by cell cycle phase.

To further test the the ability of mPOP and SCoPE-MS to quantify proteins in single cells, we sorted mouse embryonic stem (ES) cells based on the phase of their cell division cycle (CDC), Fig. 3c. To this end, we used a fluorescent protein Citrine fused to partial sequences of Geminin that is a ubiquitin-target of the anaphase promoting complex, and thus Citrine is degraded periodically during the CDC^14^. Using this system, known as FUCCI, we sorted ES from the G1 and the G2 phase of the CDC and quantified their proteomes. PCA analysis of the proteins exhibiting the gradient variation across the single cells separated these cells into clusters consistent with the CDC phase determined by the FUCCI system, Fig. 3c. Examining the proteins driving this separation, we found that, consistent with expectations, these proteins are enriched for CDC functions.

## Discussion

Minimizing sample losses and maximizing throughput is a major requirement for applying ultra-sensitive MS to biological problems^3,8^. It has motivated many colleagues to develop sample preparation methods with minimal volumes and low cleanup losses^2,6^. However, mPOP is the only method that uses solely MS-compatible reagents and allows parallel preparation of hundreds of samples. Crucially, mPOP uses inexpensive equipment accessible to most labs. Furthermore, mPOP allowed us to reduce the sample preparation volume for SCoPE-MS and reduce losses while massively increasing the throughput and the consistency of the data. Thus, mPOP empowers automated preparation of SCoPE-MS sets at much lower cost than what was possible by focused acoustic sonication. This allows to increase the number of analyzed single cell with affordable resources.

## Acknowledgments

We thank A. Petelski, R. G. Huffman, A. Chen, and J. Neveu, for assistance, discussions and constructive comments. This work was funded by startup funds from Northeastern University and a New Innovator Award from the NIGMS from the National Institutes of Health to N.S. under Award Number DP2GM123497. Funding bodies had no role in data collection, analysis, and interpretation.

## Competing Interests

The authors declare that they have no competing financial interests.

Correspondence and requests for materials should be addressed to N.S.

## Contributions

H.S., G.H, D.P., E.E., Z.N., B.B and N.S. performed experiments and collected data; N.S. and H.S designed experiments, analyzed the data and wrote the manuscript. N.S. raised funding and supervised research.

## Data Availability

The raw MS data and the search results were deposited in MassIVE (ID: MSV000082841) and in ProteomeXchange (ID: PXD010856). Supplemental website can be found at: northeastern.edu/slavovlab/mPOP/

## Methods

### Cell culture

Jurkat and U-937 cells were grown as suspension cultures in RPMI medium (HyClone 16777-145) supplemented with 10% fetal bovine serum (FBS) and 1% pen/strep. Cells were passaged when a density of 10^6^ cells/ml was reached, approximately every two days. HEK-293 were grown as adherent cultures in DMEM supplemented with 10% FBS and 1% pen/strep and passaged at 70% confluence, approximately every two days. Mouse embryonic stem cells expressing the FUCCI system were grown as adherent cultures in 10 cm plates with 10 ml Knockout DMEM media supplemented with 10 % ES certified FBS, non-essential amino acids supplements, 2 mM L-glutamine, 110 *µMβ*-mercapto-ethanol, 1 % penicillin and streptomycin, and leukemia inhibitory factor (mLIF; 1,000 U LIF/ml). ES cells were passaged every two days using StemPro Accutase on gelatin coated tissue culture plates. Starting six passages prior to harvesting, ES cells were grown in media containing DMEM/F12 with N2, B27, and NEAA supplements, 1% pen/strep, 110 *µMβ*-mercapto-ethanol, 3 mM L-glutamine, 200ug/ml human insulin, 1 *µM* PD0325901, 3 *µM* CHIR99021, and 1e4 units/ml mLIF.

### Harvesting cells for mPOP

To harvest cells, embryoid bodies were dissociated by treatment with StemPro Accutase (ThermoFisher #A1110501) and gentle pipetting. HEK-293 cells were dissociated by gently pipetting. Cell suspensions of differentiating ES cells, Jurakt cells or U-937 cells were pelleted and washed quickly with cold phosphate buffered saline (PBS) at 4 *°C*. The washed pellets were diluted in PBS at 4 *°C*. The cell density of each sample was estimated by counting at least 150 cells on a hemocytometer.

### Sorting cells by FACS

HEK-293 and U-937 cells were sorted by FACS (Beckman Coulter MoFlo Astrios EQ Cell Sorter) into 2uL of pure water in 96-well PCR plates (Eppendorf twin.tec E951020303). Mouse embryonic stem cells were sorted by FACS (BD FACSAria I) into the same type of 96-well PCR plates. The mouse embryonic stem cells express the fucci system, and were sorted based on fluorescence of the citrine-geminin fusion protein.

### Cell lysis and digestion

Bulk and single cells alike were lysed by freezing at-80 *°C* for at least 5 minutes and heating to 90 *°C* for 10 minutes. Then, samples were centrifuged briefly to collect liquid, and trypsin (Promega Trypsin Gold) and buffer triethylammonium bicarbonate (TEAB) (pH 8.5) were added to 10 *ng/µl*) and 100mM, respectively. The samples were digested for 4 hours in a thermal cycler at 37 *°C* (BioRad T100). Samples were cooled to room temperature and labeled with 1 *µl* of 43mM TMT label (TMT11 kit, ThermoFisher, Germany) for 1 hour. The unreacted TMT label in each sample was quenched with 0.5 *µl* of 0.5% hydroxylamine for 30 minutes at room temperature. Samples were centrifuged briefly following all reagent additions to collect liquid. The samples corresponding to one TMT11 plex were then mixed in a single glass HPLC vial and dried down to 10 *µl* in a speed-vacuum (Eppendorf, Germany) at 35*°C*.

### Master mix preparation

Jurkat and U-937 cells were harvested and counted as described above. Five thousand three hundred cells from each type were digested (100mM TEAB pH 8.5, 10 *ng/µl* trypsin at 37 *°C* for 4 hours), divided into 5000, 100, 100, and 100 cell equivalents, labeled with TMT11, and combined such that there are two carrier channels of 5000 cell equivalents (one of Jurkat, one of U-937) and six channels of 100 cell equivalents, three of Jurkat and three of U-937 (Fig. S1a). This sample was diluted 100x and aliquoted into glass HPLC vials. Material equivalent to 50 cells in the two carrier channels and 1 cell in the six other channels was injected for analysis by LC-MS/MS.

### Mass spectrometry analysis

SILAC data was acquired using a Dionex UltiMate 3000 UHPLC with a 25cm length × 75*µm* inner diameter microcapillary column packed with C18 Reprosil resin (1.9 *µm* resin, Dr. Maisch GmbH, Germany). Peptides were separated at 150 nL/min over a 180 minute gradient and analyzed on a Thermo Scientific Lumos mass spectrometer. After a precursor scan from 400 to 2000 m/z at 50,000 resolution the top 10 most intense multiply-charged precursors (charges 2 to 4) were selected for alternating HCD and CID fragmentation at 50,000 and 35,000 resolutions, respectively. Mouse embryonic stem cell (SCoPE-MS) data was acquired using a Proxeon Easy nLC1200 UHPLC (Thermo Scientific) at a flow rate of 200 nL/min using a 25cm length × 75*µm* Waters nanoEase column (1.7 *µm* resin, Waters PN:186008795) over a 60 minute gradient. Peptides were analyzed by a Thermo Scientific Q-Exactive mass spectrometer. After a precursor scan from 450 to 1600 m/z at 70,000 resolution, the top 5 most intense precursors with charges 2 to 4 were selected for HCD fragmentation at resolution 70,000 with a max fill time of 300ms. A 0.7 Th isolation window was used for MS2 scans.

### Analysis of raw MS data

Raw data were searched by MaxQuant^15,16^ 1.6.0.16 and 1.6.2.3 against a protein sequence database including all entries from the appropriate mouse or human SwissProt database (downloaded July 15, 2018 and July 30, 2018, respectively) and known contaminants such as human keratins and common lab contaminants. MaxQuant searches were performed using the standard work flow^17^. We specified trypsin specificity and allowed for up to two missed cleavages for peptides having from 5 to 26 amino acids. Methionine oxidation (+15.99492 Da) and protein N-terminal acetylation (+42.01056 Da) were set as a variable modifications. Carbamidomethylation was disabled as a fixed modification. All peptide-spectrum-matches (PSMs) and peptides found by MaxQuant were exported in the msms.txt and the evidence.txt files. SILAC data was searched in two batches (by date acquired) with match between runs enabled, using the default settings.

### Principle component analysis for single cell data sets

Using the data analysis language R (v3.4.1), the matrix of peptide-level quantitation from TMT reporter ions was normalized prior to PCA analysis. Columns (corresponding to separate TMT channels) were divided by their median value. Rows (corresponding to peptides from individual TMT11-plexes) were divided by their mean, then the mean of the resulting vector subtracted from all values in the vector.

### SILAC data normalization

Expected SILAC ratios for peptides were computed by taking the mean of the SILAC ratios from samples containing equal number of SILAC heavy and SILAC light U-937 cells, processed by the urea-based method. All subsequent samples, processed either by mPOP or the urea-based method, were normalized by these values to account for artifacts from SILAC labeling.

## Supplementary Figures

**Figure S1.**
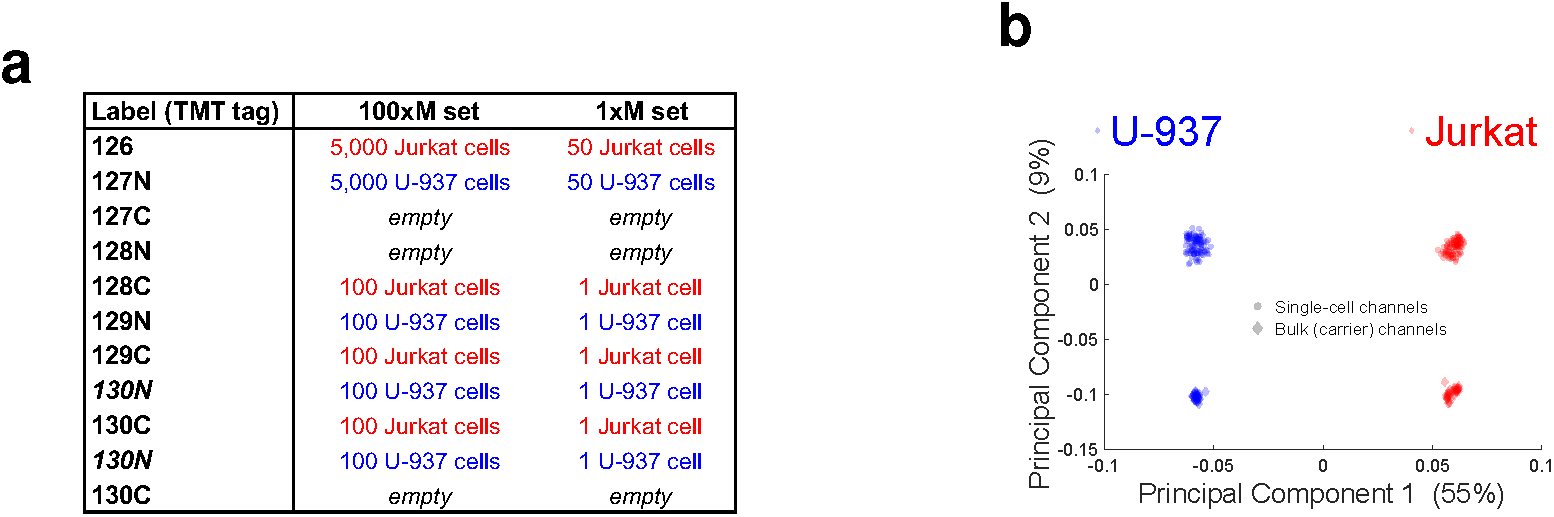
Design and quantification of 100*×M* SCoPE-MS sets. (a) Schematic for the design of 100 *× M* sets and the proteome amounts corresponding to 1 *× M* sets. (**b**) PCA of 1*×M* sets without normalization of the carrier and the single-cell channels.

